# *Lactobacillus plantarum* favors the early emergence of fit and fertile adult Drosophila upon chronic undernutrition

**DOI:** 10.1101/080549

**Authors:** Mélisandre A. Téfit, François Leulier

## Abstract

Animals are naturally surrounded by a variety of microorganisms with which they constantly interact. Among these microbes, some live closely associated with a host and form its microbiota. These communities are now extensively studied, owing to their contributions to shaping various aspects of animal physiology. One of these commensal species, *Lactobacillus plantarum*, and in particular the *L.p.*^*WJL*^ strain, has been shown to promote the growth of Drosophila larvae upon nutrient scarcity, allowing earlier metamorphosis and adult emergence compared to axenic individuals. As for many insects, conditions surrounding the post-embryonic development dictate key Drosophila adult life history traits, and adjusting developmental timing according to the environment is essential for adult fitness. The growth acceleration induced by *L.p.*^*WJL*^ occurs in a context of poor nutrition and we wondered if this could adversely impact the fitness of Drosophila adults. Here we show that the *L.p.*^*WJL*^- mediated acceleration of growth is not deleterious; adults emerging after an accelerated development are as fit as their axenic siblings. Additionally, *L.p.*^*WJL*^’s presence even leads to a lifespan extension in nutritionally challenged males. These results demonstrate that *L.p.*^*WJL*^ is a beneficial partner for *Drosophila melanogaster* through its entire life cycle. This commensal bacteria allows the earlier emergence and longer survival of fit and fertile individuals and might represent one of the factors contributing to the ecological success of Drosophila.

**Summary statement:** *Lactobacillus plantarum*^*WJL*^ is beneficial to Drosophila physiology along its entire life cycle. This bacteria triggers the early emergence and longer survival of fit and fertile adults.

## Introduction

In nature, animals are constantly surrounded by a profusion of microorganisms, whose presence has contributed to shaping life as we know it (McFall-Ngai 2015). The interactions existing between microbes and animals cover a broad spectrum, with outcomes ranging from obligate symbiosis to lethal infection (Casadevall & Pirofski 2000; Hentschel et al. 2000). Among these microbial species, some live closely associated with an animal host with which they establish commensalistic or mutualistic relationships. The community they form is referred to as the microbiota, which over the last years has been increasingly studied for its impact on various physiological traits. Indeed, in several mammalian, nematode or arthropod models, the microbiota has been shown to shape development, immunity, metabolism and even behavior (Kostic et al. 2013; Lee & Hase 2014). In this fast expanding research field, Drosophila has been a fruitful model. Thanks to its ease of manipulation and genetic tractability, as well as the low complexity of its microbiota, the fruit fly represents a powerful tool to delve into the mechanistic underpinnings of host-microbiota interactions (Lee & Brey 2012; Ma et al. 2015). Studies revealed that microbiota’s presence and composition impact various traits throughout Drosophila life cycle such as larval growth, developmental timing, stress resistance, immune response, metabolism, lifespan and behavior (Brummel et al. 2004; Ryu et al. 2008; Sharon et al. 2010; Shin et al. 2011; Guo et al. 2014; Petkau et al. 2014; Venu et al. 2014; Wong et al. 2014; Clark et al. 2015). As the microbiota is closely associated to its animal partner and, in the case of Drosophila, is an integral part of its nutritive substrate, it is not surprising to see its influence on so many biological functions. Moreover, as for many insects the larval life is a highly plastic stage in the fly life cycle. Indeed, biotic and abiotic factors surrounding the development of an organism participate in shaping this process (Gilbert 2001; McFall-Ngai 2002), and in turn have a crucial impact on several key life history traits at the adult stage, such as reproductive capacities, stress resistance or lifespan (Tu & Tatar 2003; Andersen et al. 2010; Sisodia & Singh 2012; Burns et al. 2012).

Previously, we showed that, upon mono-association, some strains of the commensal bacterial species *Lactobacillus plantarum* (a member of the dominant phyla of Drosophila’s microbiota) are able to sustain the systemic growth of Drosophila larvae to the same extent as a more complex microbiota (Storelli et al. 2011; Erkosar et al. 2015). Upon yeast deprivation during the larval stages, mono-association of germ-free animals with the strain *L.p.*^*WJL*^ isolated from the intestine of lab-raised *Drosophila melanogaster* (Ryu et al. 2008) increases larval growth and reduces developmental timing, thus allowing the earlier entry into metamorphosis of mono-associated individuals (Storelli et al. 2011).

Several studies using Drosophila lines generated in a laboratory evolution experiment of postponed senescence selection (Rose 1984) have described a series of trade-offs between key life history traits. This occurs when the optimization of a trait correlates with a negative impact on another parameter; for example increased reproductive capacities usually come at the cost of shortened lifespan. Such trade-offs can involve traits from either the same life stage or across different life stages, and thus the length of the larval period, early and late life fecundity, adult longevity as well as stress resistance were shown to trade-off with one another (reviewed in Zera & Harshman 2001). Given the numerous examples of life history trade-offs and the rather striking effect of *L.p.*^*WJL*^ on larval development, we wondered about the potential repercussions of this accelerated growth on adult fitness. We speculated that *L.p.*^*WJL*^-mediated acceleration of growth in an otherwise nutritionally challenging environment might be deleterious at later stages such that it would lead to the emergence of unfit adults. To address this question, we assessed several fitness parameters in young adult flies and observed that overall, *L.p.*^*WJL*^-association was not detrimental for adult fitness. Furthermore, for adult males it proves to be an advantageous partner; *L.p.*^*WJL*^-associated males are not only emerging several days before their germ-free siblings, they also survive longer in nutritionally challenging conditions. *L.p.*^*WJL*^ is thus a true beneficial partner for Drosophila along its entire life cycle, and even more so in a poor nutritional environment. We therefore propose that bacterial members of the fly microbiota might represent one of the factors contributing to the ecological success of *Drosophila melanogaster*.

## Material & Methods

### Fly stocks and husbandry

Wolbachia-free fly stocks (*yw*) were reared on a standard yeast/cornmeal diet containing for 1L: 50g inactivated yeast (Bio Springer, Springaline BA95/0-PW), 80g cornmeal (Westhove, Farigel maize H1), 10g agar (VWR, ref. #20768.361), 5,2g methylparaben sodium salt (MERCK, ref. #106756) and 4ml 99% propionic acid (CARLO ERBA, cref. #409553). All experimental flies were kept in incubators at 25°C, with a 12h/12h light/dark cycle. The low-yeast diets were made by decreasing the quantity of yeast to either 30, 12, 8 or 6g/L and the quantity of agar to 7,2g/L. Unless stated otherwise, only mated flies were used in this study.

### Generation of axenic Drosophila stocks and bacterial mono-association

To generate axenic flies, eggs were collected overnight and treated in sterile conditions with successive 2 minutes baths of bleach and 70% ethanol. Bleached embryos were then rinsed in sterile water for another 2 minutes and placed on sterile standard food supplemented with an antibiotic cocktail (50μg ampicillin, 50μg kanamycin, 50μg tetracyclin and 15μg erythromycin per liter of fly food). Emerging adults were tested for axenicity by crushing and plating of the fly lysate on different bacterial culture media. Germ-free flies were kept on antibiotic food for a few generations and conventionally reared stocks were used to regenerate axenic stocks regularly. For bacterial mono-association, 50μL of PBS containing 10^8^ CFU of a stationary phase culture of *Lactobacillus plantarum*^*WJL*^ were used to inoculate the surface of the food contained in a Ø1,5cm fly tube. 50 axenic eggs from an overnight collection were transferred onto the inoculated food and left to develop until adult emergence. The experimental germ-free condition was obtained by inoculating the food with sterile PBS. In case of association at the adult stage, fly food was inoculated as described above and let to dry under a hood. Forty to fifty newly emerged adult flies (females and males mixed 1:1) were then transferred into the inoculated tubes and reared for 7 days until the beginning of the experiments.

### Developmental timing

Fifty germ-free embryos were associated with *Lactobacillus plantarum*^*WJL*^ or kept axenic, as described in the previous section. Larvae were then left to develop under low nutrient conditions (low yeast diet, 8g/L yeast) and the number of pupae appearing each day was recorded until the last larvae of the population reached pupariation.

### Fecundity and fertility assessment

At emergence, groups of 5 females plus 5 males were distributed in vials and flipped every 24h in a new tube. The number of eggs laid was recorded every day for 10 days and the subsequent number of emerging adults was used to calculate the fertility ratio (number of emerging progeny/number of eggs laid). In experiments where bacterial association was done only at adult stage, the fecundity/fertility assays were started at day 7 or day 10 after adult emergence and carried on for 3 days to 7 days.

### Number of ovarioles

4 to 5 days old mated females were used to assess the number of ovarioles after development on either standard (50g/L yeast) or low-yeast (8g/L yeast) diet. Newly emerged adult flies were kept on standard food until the time of dissection. Ovaries were dissected in cold PBS and directly fixed in 4% formaldehyde for 20 minutes. They were then stained with DAPI (1:1000) for 15 minutes and transferred to 80% glycerol for preservation. After fixation and staining, ovarioles were teased apart under a dissecting microscope and mounted on slides to be counted.

### Adult wet weight and resistance to full starvation

Either virgin (0 to 7 hours old) or mature and mated adults (7 or 10 days old) were collected and pooled by 5 to be weighted on a Sartorius analytical balance CPA324S (Sartorius Weighing Technology GmbH, Goettingen, Germany). Flies of the same ages were also used for full starvation assays, in tubes providing only water supply to the flies. Specifically, the starvation tubes contain a cotton ball soaked into a water reservoir to prevent them from drying. The cotton is covered with a piece of Whatmann paper on which the flies will be. Survival of the flies was recorded twice a day until all individuals were dead.

### Lifespan

After larval development on either standard (50g/L yeast) or low-yeast (8g/L yeast) diet, newly emerged adults were kept all together for 3 to 4 days before males and females were separated for the subsequent experiments. Groups of 10 mated flies were transferred to fresh vials containing either standard or low-yeast diet. Flies were transferred to fresh fly food tubes twice a week and survival was recorded daily until all individuals were dead. Depending on the condition and on the experiment, 5 to 10 replicates were performed.

### Statistical analyses

For comparison of GF and *L.p.*^*WJL*^-associated conditions Mann-Whitney test (for weight, fecundity, fertility) and logrank test (for survival curves comparison) were performed using GraphPad Prism software version 6.0f for Macintosh (GraphPad Software, La Jolla California USA, www.graphpad.com). Whiskers of the boxplots represent the minimal to maximal values. For all experiments, the p-values were reported on the corresponding figure panels only when inferior to 0,05.

## Results

### *Lactobacillus plantarum*^*WJL*^ does not directly impact Drosophila adult fitness

To determine whether *L.p.*^*WJL*^ had an impact on the fly physiology at the adult stage, we first assessed the direct effect of *L.p.*^*WJL*^ on adult Drosophila, by associating newly emerged flies with the bacteria. After a larval development on a normal diet in axenic conditions (germ-free (GF), devoid of microbiota), the young emerging adults were either associated with *L.p.*^*WJL*^ or kept axenic **(Figure 1A)**. The flies were left to mature for several days on diets with decreasing amounts of yeast and were then tested for fecundity, fertility and resistance to full starvation. After 8 days in various nutritive conditions, there was a clear effect of the diet composition on the number of eggs laid per female and on the number of adult progeny emerging from these eggs; with decreasing amount of yeast in the diet, the flies were laying fewer eggs **(Figure 1B)** and the fertility ratio (number of emerging progeny/number of eggs laid) showed an increased variability **(Figure 1C)**. The ability of females to endure complete starvation was also impacted by the amount of yeast in the diet. Indeed, 7 days old females survived longer when they had been kept on a low-yeast diet after emergence **(Figure 1D left panel**). In contrast, the diet composition did not matter for their male counterparts, who were always dying at the same rate regardless of the diet they were kept on since emergence **(Figure 1D right panel**). The association with *L.p.*^*WJL*^ however did not impact any of these adult fitness traits. In addition, we tested the same parameters in flies that were raised on a normal diet in the presence of *L.p.*^*WJL*^ during larval life. In such optimal nutritional conditions, the developmental time is similar for the axenic and the *L.p.*^*WJL*^-associated flies, and here again there was a clear impact of the diet composition on fecundity, but no bacterial contribution was revealed for either fecundity or resistance to full starvation **(Figure S1)**. We next assayed the lifespan of these flies raised with or without *L.p.*^*WJL*^ on a normal diet, and kept as adult on either the same optimal diet or on low-yeast food **(Figure 1E,F)**. Here, we saw a significant increase in the lifespan of axenic females kept in nutritionally rich conditions throughout all their life cycle **(Figure 1G left panel**). For their male counterparts however, as well as for female and male flies that went from a larval development on a normal diet, to adult life on a low-yeast diet, there was no significant impact of *L.p.*^*WJL*^ presence **(Figure 1G right panel and 1H**). Taken together, these results show that apart from the previously described sexually dimorphic lifespan shift on a normal diet (i.e. increased lifespan in GF females; (Petkau et al. 2014; Clark et al. 2015)), association of *Drosophila* with *L.p.*^*WJL*^ does not seem to have a direct impact on the adult fitness when flies develop on a normal diet.

**Figure 1.**
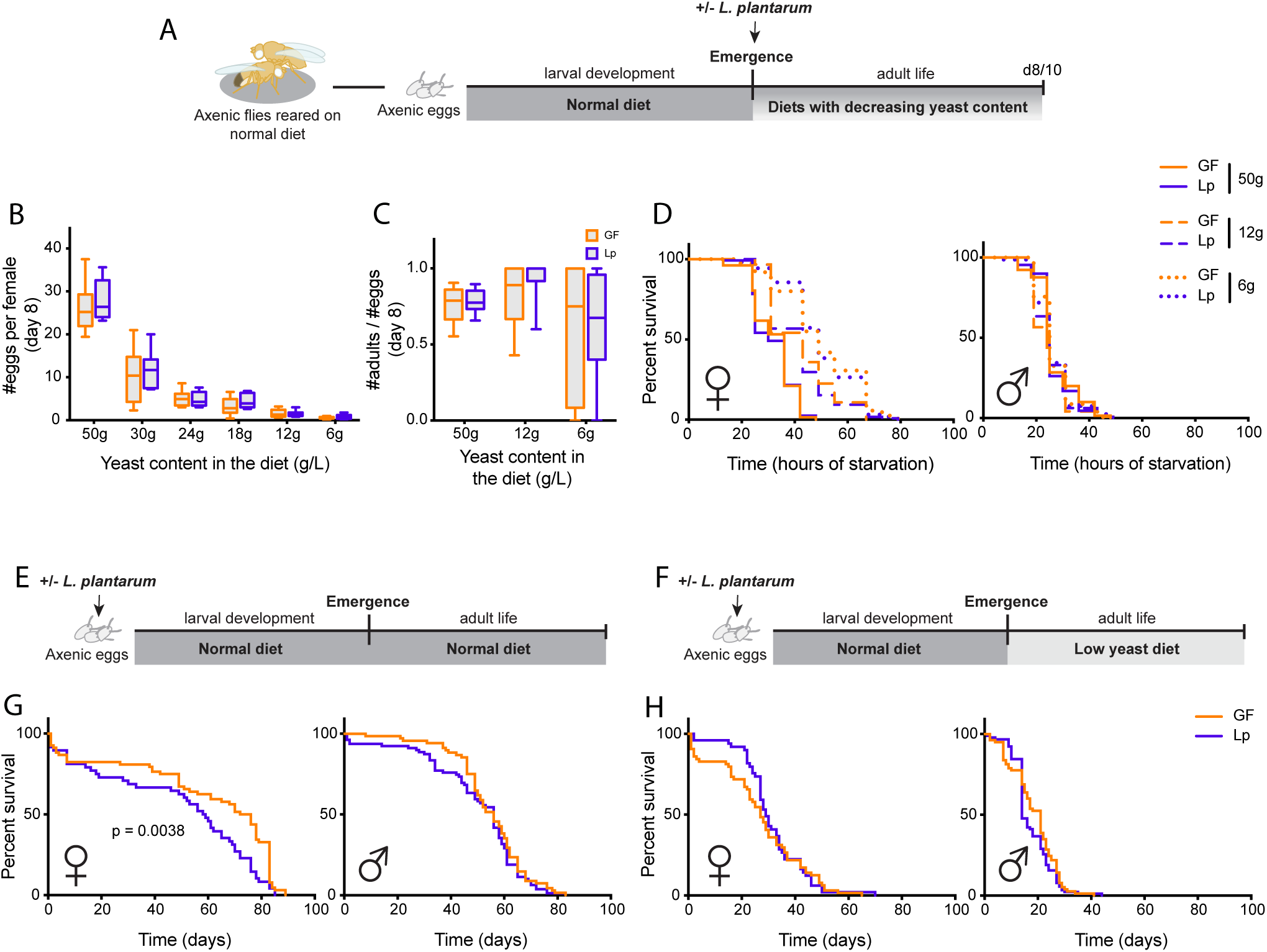
*Lactobacillus plantarum*^*WJL*^ does not directly impact Drosophila adult fitness. Right after emergence, axenic (germ-free (GF); devoid of microbiota) adults developed on a normal diet are associated with *L.p.*^*WJL*^ or sterile PBS (A). When mature, they are tested for fecundity (B) or fertility (C) at 8 days old, and for resistance to complete starvation (D) at 10 days after emergence. In panels E to H, axenic eggs were inoculated with *L.p.*^*WJL*^ or sterile PBS and developed on a normal diet (50g/L). The lifespan of the adults was then assessed, on either the same normal diet, or on a diet with a reduced amount of yeast (8g/L).

### *The Lactobacillus plantarum*^*WJL*^-mediated larval growth acceleration is not deleterious for adult fitness

While searching for a direct effect of *L.p.*^*WJL*^ on the adult stage, we did not detect any significant impact of this commensal bacteria on the tested fitness parameters. There is however a quite striking larval effect, as nutritionally challenged individuals develop faster and pupariate several days earlier when they are associated with *L.p.*^*WJL*^ compared to the axenic ones (Storelli et al. 2011; Erkosar et al. 2015). While faster larval growth and precocious emergence of the adult represent an obvious ecological advantage, doing so under nutritionally challenging conditions may in turn be deleterious for the adult fitness and reproductive success. Indeed, adjusting developmental timing to environmental cues is key to Drosophila adult fitness (Nylin & Gotthard 1998), yet upon *L.p.*^*WJL*^-association animals develop faster even though the nutritional conditions are poor. To investigate whether the growth acceleration mediated by *L.p.*^*WJL*^ upon nutrient scarcity would adversely impact subsequent adult fitness, we tested flies raised on a low yeast diet with or without the bacteria, as depicted in **Figure 2A**. As previously described, when raised on a low yeast diet larvae associated with *L.p.*^*WJL*^ pupariate several days before their axenic siblings (Storelli et al. 2011; Erkosar et al. 2015) and **Figure 2B**). We then assessed the potential repercussions of the *L.p.*^*WJL*^-association on the reproductive capacities of flies that underwent larval development in such nutritionally challenging conditions. Similar to what we observed when the flies were grown in optimal conditions and challenged only as adults, fecundity **(Figure 2F,G,H)** and fertility **(Figure 2I,J,K)** were both greatly impacted by the adult diet composition **(Figure 2C, D, E)**. The higher the yeast content in the diet, the more eggs were laid per female per day **(Figure 2F,G,H)**. In addition, the number of adult progeny emerging from these eggs was impaired on the lower yeast diet. Indeed, as we observed on the low-yeast diet in **Figure 1C** (6g/L of yeast), on the 8g/L of yeast diet the fertility ratio was greatly variable **(Figure 2I,J,K)**. On these two parameters again, there was no impact of the association with *L.p.*^*WJL*^. Furthermore, these comparable fecundity results were supported by the fact that the number of ovarioles (the functional units of Drosophila ovaries) of females raised on a low yeast diet was similar, regardless of their microbiota status **(Figure S2A)**. We are confident that our experimental setup can efficiently manipulate the ovariole number, since as expected, we observed a decreased count after a development on a low-yeast diet compared to an optimal situation (**Figure S2A** and Hodin & Riddiford 2000; Tu & Tatar 2003). As anticipated, this similar number of ovarioles between the GF and *L.p.*^*WJL*^ conditions translated into a similar cumulative number of eggs laid over the course of the experiment, and as expected we detected reduced cumulative egg laying on the poor diet **(Figure S2B)**. Next, we assayed the weight of 0 to 7 hours old virgin adults, along with their resistance to complete starvation, as readouts for direct consequences of larval life on their adult metabolic state (Baker & Thummel 2007). We detected a slight tendency in males and females associated with *L.p.*^*WJL*^ to weigh less than the axenic ones **(Figure 3A, B)**, but there was no impact of the growth acceleration mediated by *L.p.*^*WJL*^ on the flies’ ability to endure full starvation **(Figure 3C)**. These assays were repeated on mature adults, after 10 days of adult life on either a normal diet **(Figure 3D,E,F)** or on the same low-yeast diet **(Figure 3G,H,I)** and again, there was no deleterious impact of the *L.p.*^*WJL*^-mediated growth promotion on these adult fitness parameters. At this age, the weight tendency was reversed, since *L.p.*^*WJL*^-associated males and females were now slightly heavier than their axenic counterparts. Similarly to what we observed with newly emerged flies, this did not translate into differences in resistance to full starvation. In addition, similar results were obtained when we tested these parameters in adult flies that were matured on a diet with an intermediate yeast content **(Figure S3)**. Notably, for some of these experiments, the statistical analyses show significant differences between the groups, but the differences they represent are tenuous and probably not of any biological relevance. Collectively, these data suggest that even though larvae associated with *L.p.*^*WJL*^ develop faster in an otherwise poor nutritive environment, they do so without generating fitness cost for the later stage and give rise to fit and fertile adults.

**Figure 2.**
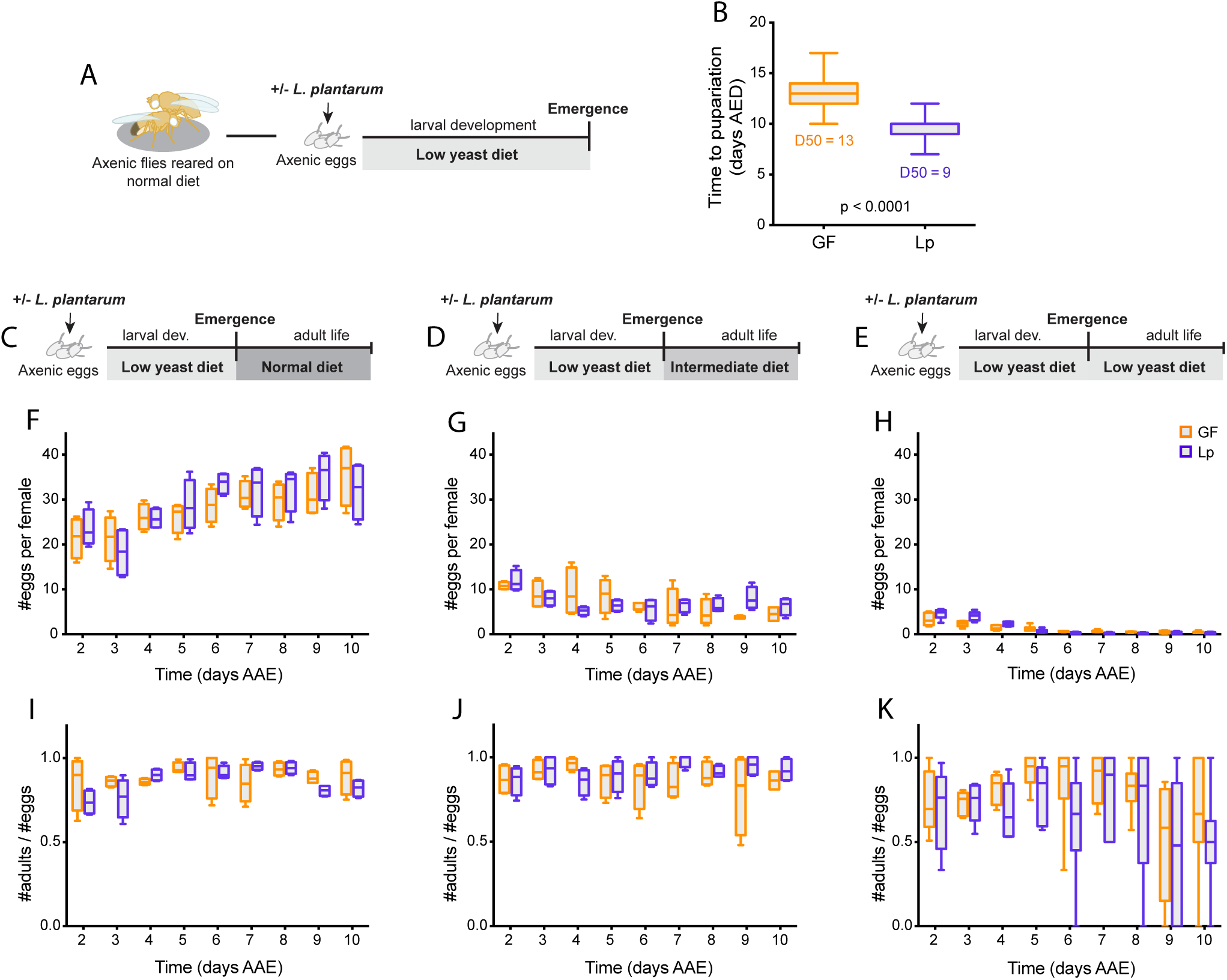
Drosophila reproductive capacities are not altered after an accelerated larval development. When larvae are raised on a low yeast diet (8g/L) (A) the presence of *L.p.*^*WJL*^ accelerates development and shortens the time to pupariation (B, days AED: after egg deposition). After this differential development adults were kept on either an optimal diet (50g yeast/L, C), an intermediate diet (30g yeast/L, D) or a nutritionally poor diet (8g yeast/L,E) and assessed for fecundity and fertility from day 2 after adult emergence (AAE) to day 10. Panels F, G and H show the number of eggs laid per female per day and the corresponding fertility ratios (emerging adult progeny divided by the number of eggs laid) are shown in panels I, J and K.

**Figure 3.**
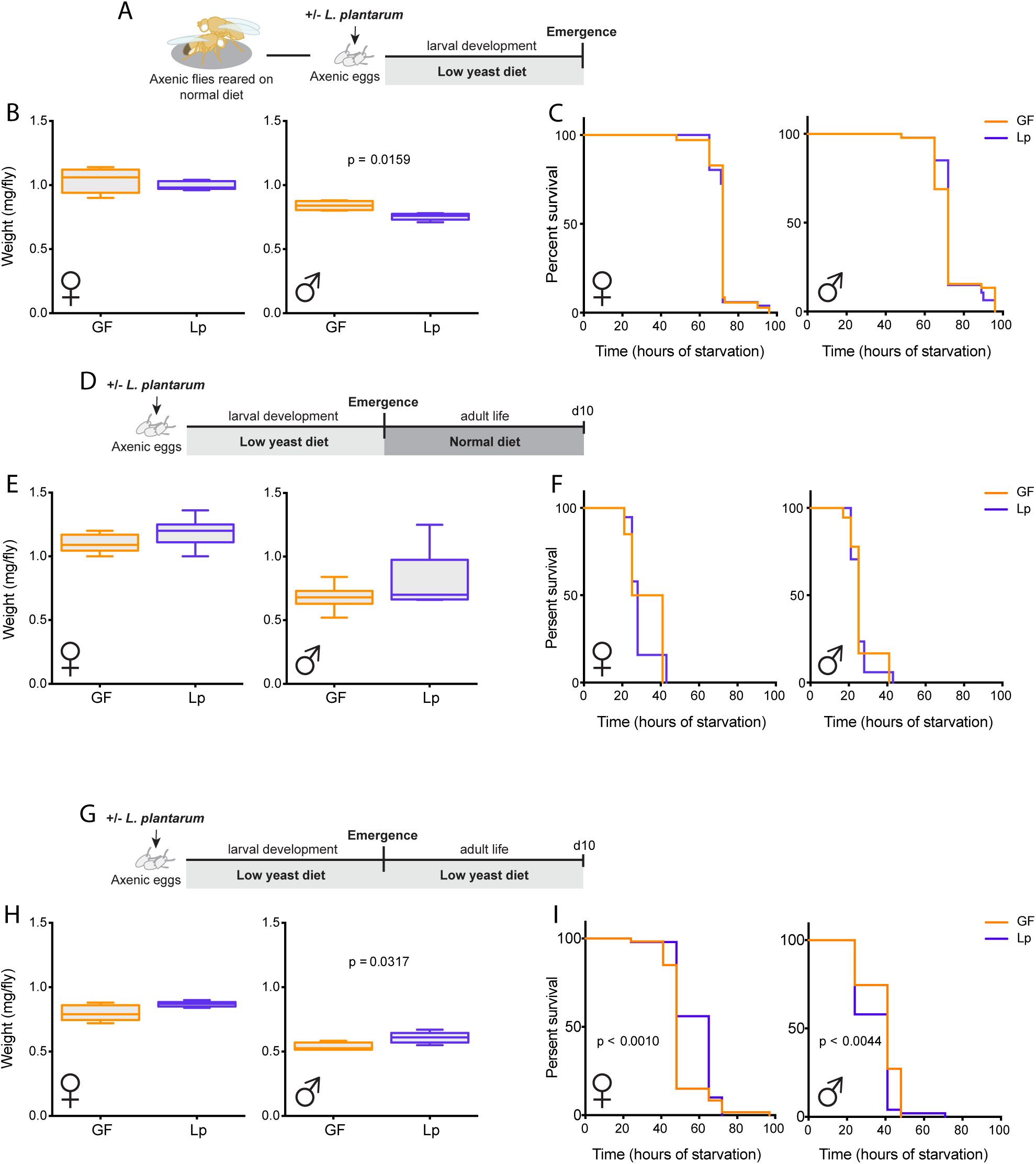
The *Lactobacillus plantarum*^*WJL*^-mediated larval growth acceleration is not deleterious for adult fitness. Right after emergence from larval development on a lowyeast diet (A), assessment of 0 to 7 hours old flies’ weight (B) and resistance to full starvation (C). The same parameters were then tested on 10 days old adults kept on either a normal diet (D-F) or on a low yeast diet (G-I).

### *Lactobacillus plantarum*^*WJL*^ increases the lifespan of nutritionally challenged males

While performing the experiments, we noticed that when kept on a low-yeast diet, adult males were dying rapidly and a significant proportion of them were already dead 10 days after emergence. We then decided to study more in details the lifespan of flies raised in such nutritionally poor conditions. After emergence from larval development on a low-yeast diet, the adults were either kept on the same low-yeast diet **(Figure 4C)**, or transferred to an optimal diet **(Figure 4A)**. We saw that, while the association with *L.p.*^*WJL*^ did not impact the lifespan of adults kept on a normal diet **(Figure 4B)**, males maintained on a low-yeast diet throughout their entire life survived better when they were associated with *L.p.*^*WJL*^ **(Figure 4D right panel and Figure S4**). Notably, their median lifespan was extended by 4 to 16 days, depending on the experiment. This fluctuation in the actual day count across experiments is commonly seen in lifespan studies (Blanc et al. 2013) but the trend persisted and was statistically significant. This result shows that in a nutritionally challenging environment, *L.p.*^*WJL*^-association not only shortens Drosophila developmental time, it also increases significantly the lifespan of adult males.

**Figure 4.**
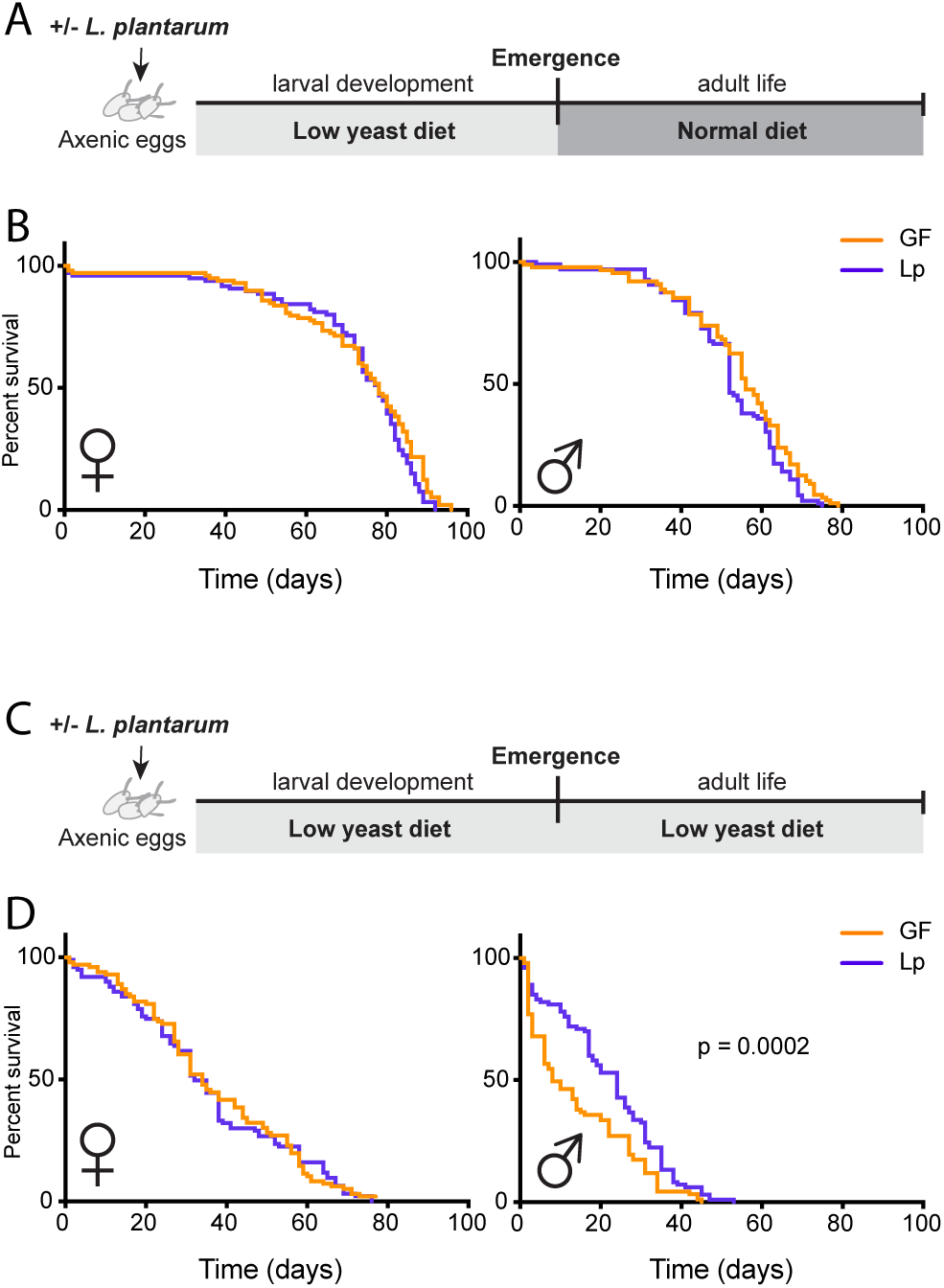
*Lactobacillus plantarum.*^*WJL*^ increases the lifespan of nutritionally challenged males. After larval development under low nutrition with or without *L.p.*^*WJL*^, the lifespan of adult males and females was assessed, when they were kept on either a normal diet (A, B) or the same low yeast diet as the larvae (C, D).

## Discussion

The microbiota is one of the key environmental factors impacting animal development and physiology and has been increasingly studied over the last few years (Sommer & Bäckhed 2013). Our work focuses on the association between *Drosophila melanogaster* and one of its natural commensal partners, *Lactobacillus plantarum*^*WJL*^. In this study we broaden our understanding of the relationship between these two partners and show that *L.p.*^*WJL*^ is beneficial for the fly throughout all life stages. We first tested if *L.p.*^*WJL*^ had a direct impact on adult fitness traits, by associating newly emerging flies after a larval development in axenic conditions. We show that, in our setup, the bacterial presence is completely dispensable for the adult fitness; the flies lay the same amount of eggs and resist starvation equally well, whether they are mono-associated or not. Notably, and contrary to *L.p.*^*WJL*^, the composition of the diet markedly impacts these parameters, and we observe a negative correlation between starvation resistance and egg laying. When decreasing the amount of yeast in the diet, we see an extension of survival upon complete starvation for female flies, together with a drop in the number of eggs laid. This effect is not only microbiota-independent but also sex-specific and the starvation resistance of males was not impacted by the quantity of yeast in the diet. Similar observations were previously reported in a study by Chippindale and colleagues, where starvation resistance is promoted by lower yeast levels in the diet, at the expense of fecundity (Chippindale et al. 1993). As in the present study, this effect was restricted to females, a characteristic that the authors attributed to distinct lipid requirements between the sexes.

In a previous study, we compared the transcriptomes of germ-free versus poly-associated flies that had been inoculated at adult stage with a cocktail of four bacterial species selected to represent the main commensals of Drosophila (*Acetobacter pomorum, Commensalibacter intestini, Lactobacillus brevis* and *Lactobacillus plantarum*). This analysis revealed a differential expression of several genes pertaining to metabolic processes; out of 105 transcripts upregulated upon bacterial poly-association, 74 were metabolism-related (Erkosar et al. 2014). With such a differential expression of metabolic genes, one could expect that certain fitness parameters, like reproductive capacities or starvation resistance, would be affected. The fact that our present study revealed no differences between germ-free and *L.p.*^*WJL*^-associated animals for these traits might be attributed to the association set-up. In this study, the flies are mono-associated with one species of *Lactobacillus* while in Erkosar et al. the animals were poly-associated. Moreover, the bacterial cocktail used in Erkosar et al. contained a species belonging to the *Acetobacter* genus. *Lactobacilliaceae* and *Acetobacteraceae* bacteria are the most represented in the communities associated with Drosophila populations, in laboratory stocks as well as in wild-caught flies (Broderick & Lemaitre 2012; Staubach et al. 2013; Chaston et al. 2016). Several studies have shown the impact of *Acetobacter* species, notably *A. pomorum* and *A. tropicalis*, on the metabolism of adult Drosophila, both upon mono-association with one species or in bacterial mixture (Newell & Douglas 2014; Huang & Douglas 2015; Chaston et al. 2016; Elgart et al. 2016). Furthermore, these studies were also specifically addressing the differential impact of *Acetobacter* species versus *Lactobacillus* species and demonstrated that, in their setup, the later had little to no effect in comparison to the former (Newell & Douglas 2014; Huang & Douglas 2015; Chaston et al. 2016; Elgart et al. 2016). It must be pointed out however, that beyond the distinction between bacterial species, the strain considered is important. Indeed, our lab and others have shown that various microbial effects are strain-specific (Storelli et al. 2011; Chaston et al. 2014). Nevertheless, taken together all these observations suggest that adult Drosophila fitness traits might be influenced by the presence of *Acetobacter* species rather than *Lactobacilli*.

After having ruled out a direct impact of *L.p.*^*WJL*^ on adult fitness, we wanted to investigate the potential repercussions of the bacteria-mediated larval growth acceleration on the adult flies. When larvae are raised on a low-yeast diet, the presence of *L.p.*^*WJL*^ promotes their growth and shortens their developmental timing (Storelli et al. 2011; Erkosar et al. 2015 and this study). However, numerous studies have demonstrated that conditions impacting larval development are known to affect several adult traits in Drosophila and a shorter larval period could negatively trade-off with adult reproductive capacities, stress resistance or longevity (Zera & Harshman 2001). We therefore suspected that this increased growth rate upon nutritional challenge could in turn adversely impact adult fitness. Here, we demonstrate that *L.p.*^*WJL*^-associated individuals are as fit as their germ-free siblings; they show similar reproductive capacities and resist complete starvation equally well, regardless of their developmental history. The association with *L.p.*^*WJL*^ is thus overall profitable to the fly, since it promotes larval growth and the early emergence of the imago without impairing the fitness of this mature and reproductive stage.

Strikingly, we find that *L.p.*^*WJL*^ extends the lifespan of males kept on poor nutritive conditions. Males that have been kept on a low-yeast diet throughout their entire life cycle benefit from the bacterial presence both as larvae and as adults; they displayed a shortened developmental timing as well as an increased median lifespan compared to their germ-free siblings. Thus, *L.p.*^*WJL*^-associated males are not only developing faster and emerging several days before their axenic counterparts, they also survive longer. In the wild, where nutrients can be scarce, longer lifespan could grant these individuals with more opportunities to mate, and to produce potentially more numerous progeny. However, to confirm this hypothesis, it is imperative to prove that these early-emerged and long-lived males are superior in their healthspan. In this light, it might be of interest to assay the late-life reproductive capacities of *L.p.*^*WJL*^-associated versus germ-free flies and see if, in addition to confer them the ability to live longer, *L.p.*^*WJL*^ also allows males to stay fit and reproductively active longer. This is an interesting future direction to follow given the growing evidences supporting a role of the microbiota in the aging process (Heintz & Mair 2014).

Overall, our results reveal that *L.p.*^*WJL*^ is beneficial for *Drosophila melanogaster* all along the fly life cycle. Indeed, upon nutritional challenge the bacterial presence allows the earlier emergence of fit and fertile adults and, in certain conditions, it even increases the lifespan of males. This *Lactobacillus* strain thus represents an advantageous partner for the fly, and taken together our results indicate that commensal bacteria might be one of the factors contributing to the ecological success of Drosophila.

## Acknowledgments

The authors would like to thank the Arthro-Tools platform of the SFR Biosciences (UMS3444/US8) for providing Drosophila husbandry materials, Loan Bozonnet for fly food preparation and Antonin Crumière, Claire-Emmanuelle Indelicato, Renata Matos and Dali Ma for occasional help in the lifespan experiments follow up. We also thank Dali Ma for proofreading of the manuscript.

## Authors contribution

FL supervised the work. MT and FL designed the experiments. MT performed the experiments. MT and FL analyzed the results. MT wrote the manuscript with inputs from FL.

